# Lamina-specific neuronal properties promote robust, stable signal propagation in feedforward networks

**DOI:** 10.1101/596676

**Authors:** Dongqi Han, Erik De Schutter, Sungho Hong

## Abstract

Feedforward networks (FFN) are ubiquitous structures in neural systems and have been studied to understand mechanisms of reliable signal and information transmission. In many FFNs, neurons in one layer have intrinsic properties that are distinct from those in their pre-/postsynaptic layers, but how this affects network-level information processing remains unexplored. Here we show that layer-to-layer heterogeneity arising from lamina-specific cellular properties facilitates signal and information transmission in FFNs. Specifically, we found that signal transformations, made by neighboring layers of neurons on an input-driven spike signal, are complementary to each other. This mechanism boosts information transfer carried by a propagating spike signal, and thereby supports reliable spike signal and information transmission in a deep FFN. Our study suggests that distinct cell types in neural circuits have complementary computational functions and facilitate information processing on the whole.

**Significance Statement:** Neural systems have many cell types that differ in properties such as size, shape, cellular mechanisms, etc. Furthermore, neurons often propagate signals to other neurons that have properties very different from their own. We investigated what this phenomenon implies in neural information processing by using computational network models, inspired by a recent experimental study on the olfactory neural pathway of fruit flies. We found that different types of neurons can perform complementary functions in a network, which boosts information transfer on the whole and supports robust, stable signal propagation in a “deep” network with many layers. Our study demonstrates that diverse cell types with different intrinsic properties are crucial for optimal signal and information transfer in neural networks.

## Introduction

How different cell types in a neural system contribute to signal processing by a whole circuit is a prime question in neuroscience. Experimental investigations of this question are increasingly common, especially due to advances in observing and manipulating neurons with particular genetic signatures. Feedforward circuits are notable targets of those studies, since, in many systems, they have been observed to comprise cell groups or “layers” with properties distinct from those of other layers, in size, morphology, expression of membrane/intracellular mechanisms, etc. For example, in the *Drosophila* antennal lobe (AL), projection neurons (PN) tend to show noisy firing, slow responses to the onset of olfactory receptor neuron (ORN) firing, and static voltage thresholds for spike generation, whereas postsynaptic neurons of PNs in lateral horns (LHN) are less noisy, fire early, and have dynamic firing thresholds (1). Also, in the cerebellum, the granule cells are tiny neurons with a simple morphology, but their postsynaptic targets, Purkinje cells, are large, with complex dendritic trees. In primary sensory cortices, the spiny stellate neurons in layer IV express NMDA receptors with the *NR2C* subunit whereas their feedforward targets, pyramidal cells in layer II/III, do not (2). These phenomena raise questions about the role of intrinsic properties and their laminar specificity. However, most theoretical and computational studies rarely take into account neuronal heterogeneity.

We addressed this question by studying the classical problem of how a spike signal, defined by evoked firing of multiple neurons in one layer, can stably propagate through multiple downstream layers in an FFN (3-11). Stable propagation plays a key role in models of conscious perception (11, 12), short-term memory (13), learning in deep artificial networks(14), etc. Most of those studies assumed that FFNs have identical types of neurons, and thus each layer makes similar input/output transformations. In this case, an input-driven spike signal either gets stronger or weaker as it passes through layers, depending on the efficacy of output spike generation, given input spikes, and also given the characteristics of the network input (Fig. 1A, Left). Then, the signal eventually reaches a fixed point of layer-to-layer transformation or dissipates (3, 5, 10) (Fig. 1A, Right). In this scenario, stable signal transmission is achieved by certain conditions for a non-trivial fixed point, which are often not robust for a wide range of initial signals. Also, irreversible signal distortion during propagation can cause inevitable loss of information.

**Figure 1.**
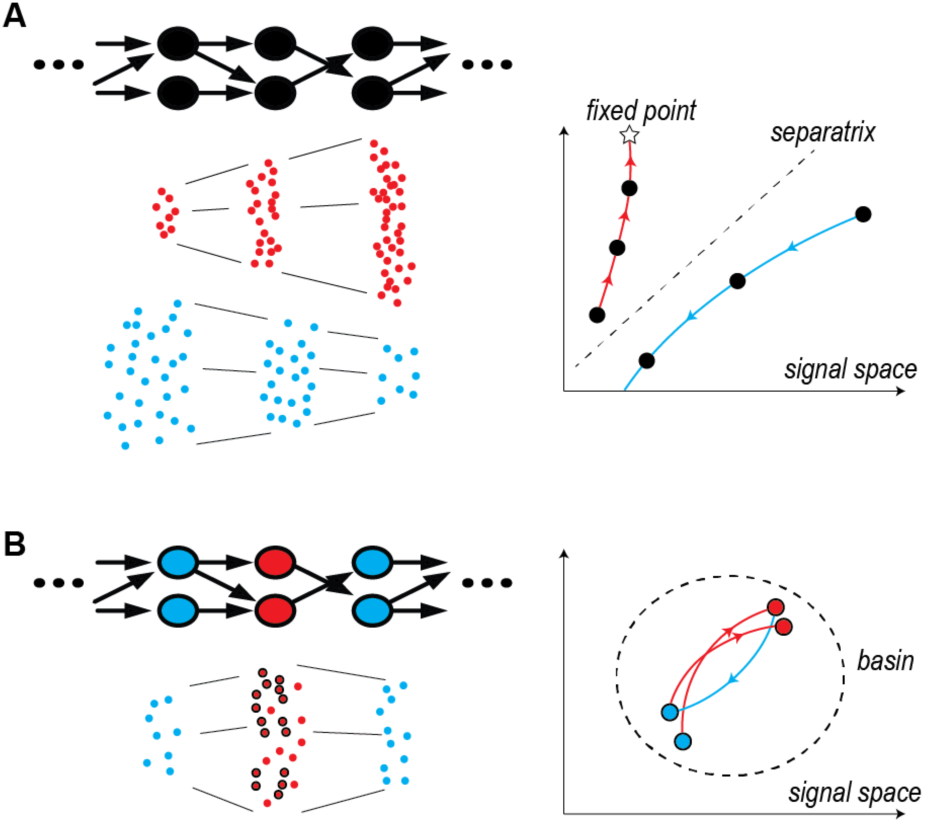
Lamina-specific intrinsic properties enable neurons to perform complementary computations in a neural network. **A.** Left: FFN with a single cell type (Top), and spikes at each layer, in two different modes of signal propagation (Bottom). One mode is amplification by progressively evoking more and more synchronized spikes (red dots) and the other is dissipation by gradually losing spikes (blue dots). Right: Trajectories of the two propagating signals, in a signal space. The x-and y-axes represent independent signal characteristics, such as the number of spikes, temporal precision, etc. A star is a fixed point of neuronal signal transformation, and a dotted line is a separatrix separating the two modes. **B.** Left: FFN where neurons have lamina-specific intrinsic properties (Top). Each layer performs a “complementary” transformation, and can selectively transfer a subset of input spikes (circled red dots), ignoring those that cause signal distortion (Bottom). Right: Trajectory of a propagating signal in a signal space. The dotted circle surrounds a region (basin) where the propagating signal is confined by the complementary transformations.

Introducing lamina-specific intrinsic properties in neurons can change this fundamentally (Fig. 1B). If each layer transforms a propagating signal in a different direction than the previous one, a fixed point will not exist in general. Instead, this prevents repeated transformation of the signal in one direction and the overall signal distortion over multiple layers can actually become smaller, compared to networks with identical layers. In particular, if transformations are “complementary,” i.e., the transformation by one layer is in the opposite or nearly opposite direction to those by its presynaptic layer, stable propagation is possible with bounded signal distortion across multiple layers (Fig. 1B Right). This mechanism can also improve information transfer. Signal distortion at each layer will accumulate if neurons in each layer repeatedly encode similar preferred features of a network input and this will cause irrevocable loss of information. Contrarily, in the case of complementary transformations, postsynaptic neurons have preferred features distinct from those of presynaptic neurons. Postsynaptic neurons filter presynaptic output spikes more selectively, resulting in demodulation of distortions introduced by presynaptic signal transformation (Fig. 1B Left). In this manner, FFNs with heterogeneous, lamina-specific neuronal properties can show enhanced information transmission compared to homogeneous FFNs.

Here we demonstrate how complementarity and robust, stable signal transmission arise from laminar specificity of cell intrinsic properties by computational FFN models. We first introduce a model of the *Drosophila* AL network with three layers of ORNs, PNs and LHNs. We show that this model replicates a recent experimental finding that differences in spiking dynamics between PNs and LHNs can balance accuracy and speed in processing olfactory information (1), and furthermore demonstrate that PN-to-LHN information transfer is nearly optimal. Then, we extend the model to a deep FFN and demonstrate robust and stable spike signal propagation, contrary to models with no laminar specificity in neuronal properties.

## Results

### Voltage-sensitive K+ channels control dynamical input/output properties of neurons

We constructed a computational model of the *Drosophila* AL network with lamina-specific neuronal properties with conductance-based neuron models, containing voltage-dependent Na^+^ and K^+^ channels (15). An important parameter of the model is *β*_*w*_, the half-activation voltage of the K^+^ channel (see Equation 1 in **Materials and Methods**). With lower *β*_*w*_, the channel is more active at subthreshold voltages, shifting the balance between inward and outward currents around the firing threshold. This fundamentally changes the neuronal signal transformation property by strengthening differentiator-like traits, whereas higher *β*_*w*_ promotes integrator-like behavior (15, 16). For example, typical repetitive firing with a sustained current input, seen in neurons with high *β*_*w*_ (=5 mV in our model), was suppressed in those with low *β*_*w*_ (=-19 mV in our model), whereas the low *β*_*w*_ case showed robust sensitivity to the dynamic fluctuation in inputs, demonstrated by evoked firing (Fig. 2A). Therefore, we will call our model neurons with low *β*_*w*_ and high *β*_*w*_ differentiators and integrators, respectively.

**Figure 2.**
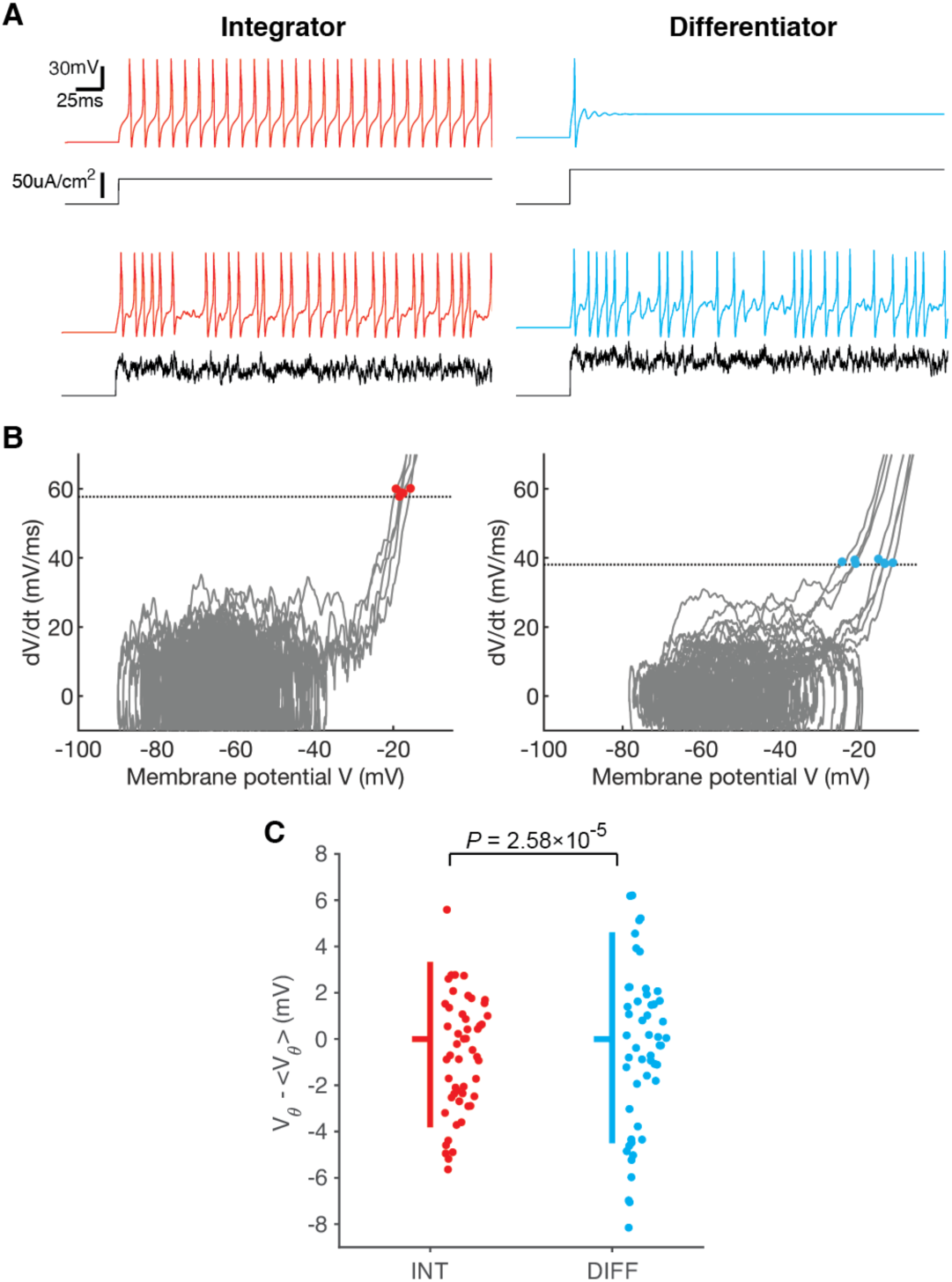
Intrinsic properties of conductance-based model neurons control dynamicity of spiking thresholds. **A.** Membrane potential response (color) to constant or fluctuating current injection (black). **B.** Example membrane potential *V* vs. *dV/dt* in two neurons, based on simulation data in Fig. 3A,B. Data from one trial are shown (gray). Dotted lines represent [*dV/dt*]_min_, the minimal *dV/dt* for spiking, and colored dots are threshold-crossing points. **C.** Spread of membrane potentials at crossing points, *V*_*θ*_, from the average. Vertical bars span from 10% to 90% quantiles, and notches are at medians. Data are the same as B, and only 50 samples (dots) are shown for clarity.

Differentiator neurons are also known to have a dynamic spiking threshold, which was also observed in LHNs (1). With the dynamic threshold, their firing depends not only on the membrane potential, but also on its temporal change, which is crucial to their sensitivity to input fluctuations (15, 17, 18). To demonstrate this, we estimated a minimal rate of membrane potential change, [*dV/dt*]_min_, that preceded spikes, but not subthreshold fluctuations from our simulation data, which corresponds to the minimal inward current required for spiking (Fig. 2B). Then, the threshold voltage, *V*_*θ*_, at *dV/dt*≈[*dV/dt*]_min_, was significantly more distributed in differentiators (Integrator: STD[*V*_*θ*_]=2.76±0.08 mV, Differentiator: 3.33±0.11 mV; *P*=2.58×10^−5^, *F*-test; Fig. 2C). This shows that differentiators can generate enough inward current to generate a spike across a broader range of membrane voltages than integrators, which is an indication of a more dynamic spiking threshold. Therefore, we used differentiator neurons, with the low-threshold K^+^ channel, for modeling LHNs, and integrators with the high-threshold channel for PNs in the AL network.

### Lamina-specific neuronal properties are crucial for the Drosophila AL network

Our network model of integrator PNs and differentiator LHNs (see **Materials and Methods** and **Table S1-S2** for full description) reproduced the key features of experimental results in ref. (1). When ORNs were given a common current input that simulates optogenetic stimulation in experiments (Fig. 3A), PNs showed a slower amplified response to transient inputs from ORNs, and LHN firing was more temporally refined, with the peak of their firing rate preceding that of presynaptic PNs, just as in experimental data (Fig. 3B Left). This rapid response of LHNs caused detection accuracy (*d*’) (see ref. (1) and **Materials and Methods**) for the ORN input to grow much faster to a larger maximum in LHNs than in PNs (Fig. 3B Right). In contrast, homogeneous networks, in which PNs and LHNs are of the same type, showed suboptimal behaviors, such as delayed firing of LHNs; therefore, *d*’ of LHN rose more slowly and reached a lower maximum than that of PNs (Fig. 3C).

**Figure 3.**
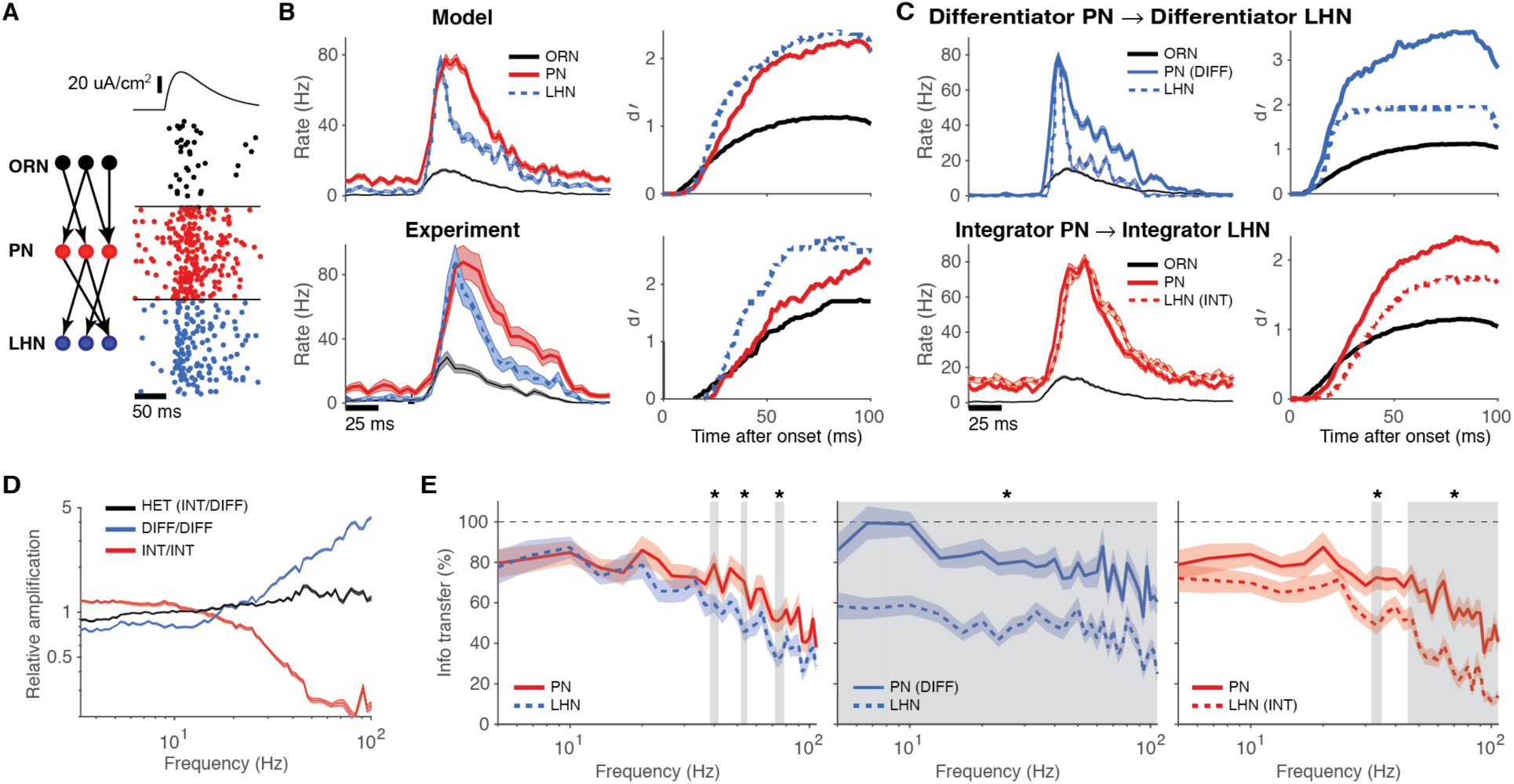
Lamina-specific neuronal properties boost information transfer in the AL network. **A.** Schematic diagram of the network model (Left), and spikes from a simulation with a current input to ORNs on top (Right). 40 trials are shown for one example neuron in each layer. **B.** Left: Average firing rates from the simulation (Top) and experimental data (Bottom). Right: *d*’ for detecting an input to ORNs at each layer, computed from the same data as Left. **C.** Same plots as B, with a model with differentiator PNs (Top), and with integrator LHNs (Bottom). **D.** Spectral power amplification, *P*_*LHN*_(ω)/*P*_*ORN*_(ω), normalized by total power. *P*(ω) is a power spectral density of mean firing rate. Black represent the heterogeneous network, while blue and red are homogeneous ones with differentiator and integrator PN/LHN, respectively. **E.** Information transfer from ORNs to PNs (solid) and LHNs (dotted). Black dotted lines represent 100% information transfer. Grey regions and stars represent frequency bands with significant differences between PNs and LHNs (*:*P*<0.01, Student *t-* test). Data are mean±SEM.

How do the different intrinsic properties of neurons contribute to the speed and high fidelity of LHN output? Since PNs and LHNs have opposite traits of differentiators and integrators, respectively, their effects can compensate for each other in the combined feedforward transformation of the ORN output. To analyze how the PN and LHN layers transform ORN inputs together, we computed how they amplify the power spectrum of ORN firing within a physiological frequency band (≲100 Hz) with data from longer simulations with continuous current stimulus to ORNs (see **Materials and Methods**). This showed that homogeneous networks with differentiator and integrator PNs/LHNs preferentially amplified higher or lower frequency components, respectively, whereas the heterogeneous network showed little distortion across the entire frequency range, demonstrating that PNs and LHNs complemented each other (Fig. 3D).

We found that this complementarity also facilitated information transfer. We estimated (the lower bound of) mutual information (MI) between the input to ORNs and spike outputs of each layer, and compared how much information in ORN firings pertaining to the input is transmitted to output firing of PNs and LHNs. Specifically, we measured the information transfer from ORNs to PNs or to LHNs by computing a ratio of MIs, *I*(ORN input; PN or LHN output)/*I*(ORN input; ORN output), respectively, where *I*(*X*; *Y*) denotes MI between *X* and *Y*. We found that information transfer to PNs closely matched that to LHNs in the heterogeneous network, whereas significant information loss was observed in homogeneous networks (Fig. 3E). In particular, the all-integrator PN/LHN case showed information loss specifically in the high frequency band (Fig. 3E Right), indicating that large signal distortion in this regime (Fig. 3D) impaired information transfer. This suggests that complementarity enabled nearly optimal information transfer from PNs to LHNs, demonstrating the importance of laminar specificity of intrinsic and functional properties of neurons.

### Lamina-specific neuronal properties promote robust and stable signal propagation in deep FFNs

We then investigated whether this mechanism can also enhance signal transmission in larger networks. For this purpose, we extended the AL network to a deep heterogeneous FFN model, by adding more alternating layers of integrator or differentiator neurons (see **Materials and Methods** and **Table S3** for full description). Then, we simulated how a packet of spikes, injected into the input layer, propagates through subsequent layers (3-11).

We found that the spike signals stably propagated in this network, whereas homogeneous networks, with only differentiators or integrators, showed opposing results (Fig. 4A,B): In the all-differentiator network, the evoked spike signal became increasingly synchronized and propagated as layer-wide synchronized spikes, whereas in the all-integrator network, the evoked spike signal became broader and less synchronized, until it was eventually lost among spontaneously firing spikes (Fig. 4A Right). Stable propagation in the heterogeneous network was decidedly robust over a wide range of input signals with diverse temporal width (σ) and total number of spikes (α) (Fig. 4C Top). Conversely, the all-differentiator network exhibited clear preference for sharply synchronized spikes (3), while signals gradually dissipated into spontaneous activity in the all-integrator network (Fig. 4C Middle-Bottom). Therefore, when tested with input signals with diverse (σ, α), the heterogeneous network showed the best performance in signal propagation (Fig. S1), and this result did not significantly change with additional feedforward inhibition in the deep FFN (Fig. S2).

**Figure 4.**
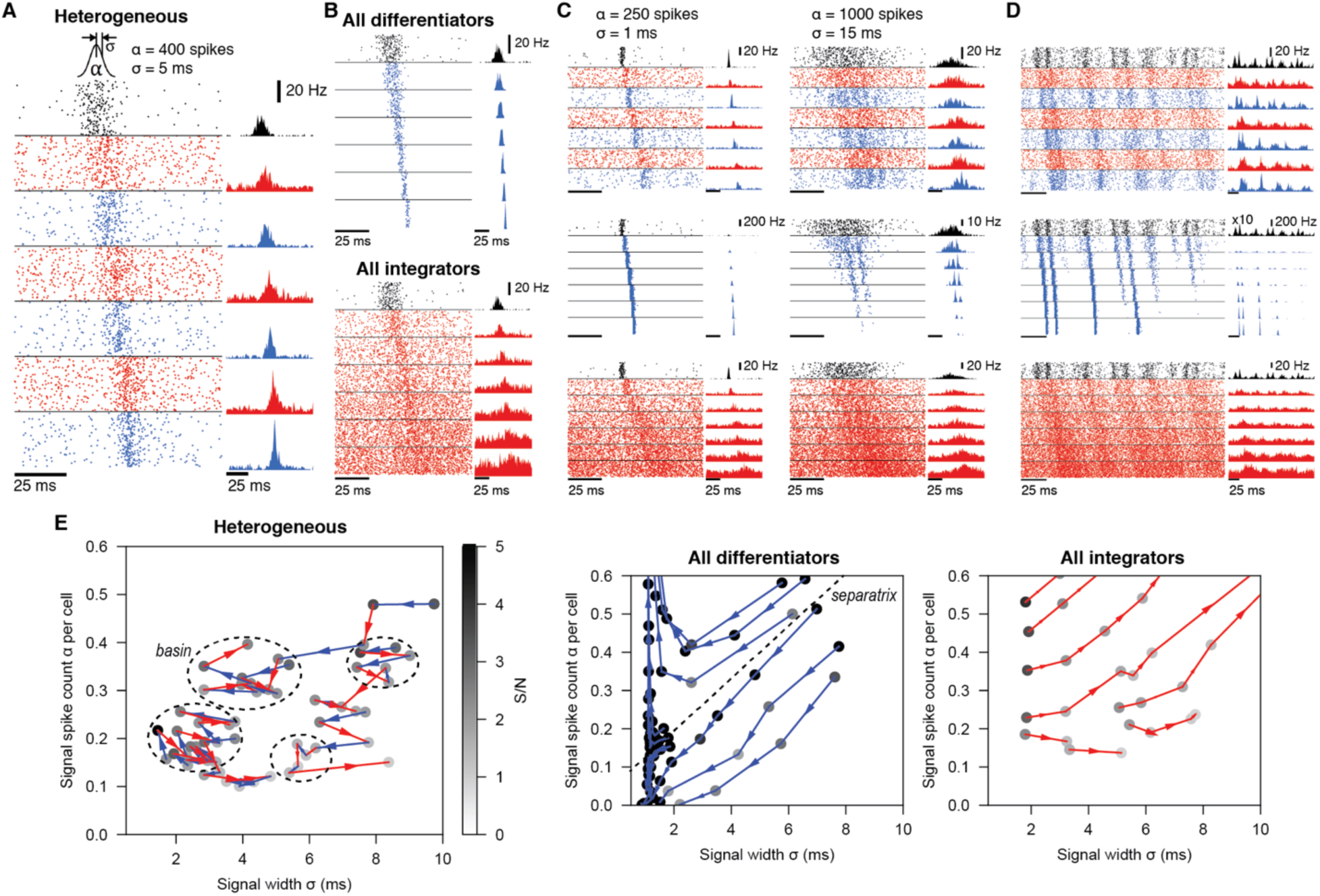
Lamina-specific neuronal properties robustly stabilize spike signal propagation in deep FFNs. **A.** Propagation in a heterogeneous network. Inset on top is the Gaussian distribution of spikes, evoked in the input layers (black). Dots (Left) and histograms (Right) are spikes and their firing rates, respectively. In all figures, blue and red represent differentiators and integrators, respectively.**B.** Firing in homogeneous networks with only differentiators (Top) and integrators (Bottom). **C.** Propagation of signals with different spike count (α) and width (σ). **D.** Network firing with continuous noise current in the input layer. In the middle row (blue; all differentiator), the input layer firing rate is multiplied by 10 for clarity. **E.** Analysis of signal transformations underlying stable propagation in the (σ, α) space. Each trajectory is formed by connecting (σ, α) of a propagating signal (dots) between adjacent layers, starting from the second layer output. Shade of each dot is the signal-to-noise ratio (S/N) and only points with S/N>1 are shown. Dotted circles mark “basins” (Fig. 1A) where any propagating signal stays for ≥5 layers. A dotted line in the Middle panel is an approximated separatrix between trajectories toward a fixed point and dissipation (Fig. 1A). All models have 9 layers and the first 7 layers are shown in A-D for clarity.

Furthermore, when the input layer fired with dynamically varying σ and α, due to dynamical, stochastic current injection, this continuous signal propagated in the heterogeneous network with many conserved features, whereas significant signal distortion and loss were again observed in the homogeneous networks (Fig. 4D). Note that propagation of dynamical input features indicates superior information transfer in a heterogeneous network, compared to homogeneous ones.

Again, complementary transformations by neighboring layers with distinct neuron types underlie the robust and stable signal propagation. To demonstrate this, we analyzed trajectories of propagating signals in the (σ, α) plane (3, 10) (Fig. 4E), a simple version of the signal space that we previously discussed (Fig. 1). In the heterogeneous network, each layer transformed an incoming signal into a different, sometimes nearly opposite or complementary direction in the (σ, α) plane than those transformed by its pre-and postsynaptic layer, which prevents formation of a uniform flow. This prevents a propagating signal from running away and confines it to a small region (basin), corresponding to stable propagation (Fig. 4E Left). In contrast, in homogeneous networks, all layers perform similar transformations and drive propagating signals rapidly toward a fixed point of sharp synchronization or dissipation (Fig. 4E Middle, Right). Notably, in most of the (σ, α) plane, transformations in those two networks are in nearly opposite directions: In the all-differentiator network, σ and α both tend to decrease (Fig. 4E Middle), because sharply correlated spikes are the preferred input of the neurons, while σ and α increase in the all-integrator network (Fig. 4E Right). In the heterogeneous network, those two different transformations are performed by neighboring layers, so as to complement each other, minimizing overall signal distortion and boosting information transfer. In summary, complementary transformations by different neuron types can protect a propagating signal from undergoing a loss or distortion regime in the signal space, instead supporting its robust and stable transmission.

## Discussion

Diversity of cell types is one of the distinctive characteristics in neural systems and its functional characterization is the subject of ongoing experimental investigations. Integrating information about cell types and their intrinsic properties with network connectivity should be an important research question to develop a holistic understanding of how spike signals propagate in neural circuits. However, diversity of cellular properties is one of the most neglected elements in theoretical neural network studies. Here we showed using our computational FFN models with various types of excitatory neurons that different cell types are beneficial in neural networks, because their different input/output transformation properties can complement each other, enhancing signal and information transmission in the whole network.

We focused on functionally distinct cell types due to different voltage-dependencies of K^+^ channels, which can arise from diverse expression patterns of low-threshold K^+^ channels (19-21). However, other neuronal mechanisms that affect the integrative cellular property can play similar roles, such as morphology (22), a high conductance state (23), inactivation of Na^+^ channels (24, 25), h-channels (26), etc. Furthermore, synaptic and circuit mechanisms known to operate as integrators or differentiators can be organized by a similar principle, such as short-term synaptic depression and facilitation, which can act as high-or low-pass filters, respectively (27), and inhibition, which can limit an integration time window for incoming inputs and promote temporal fidelity of neuronal responses (28). Our complementarity hypothesis predicts that integrator neurons, such as PNs, tend to have synapses with short-term depression (29) whereas differentiators, such as LHNs, have facilitating synapses.

Jeanne and Wilson compared spike signal transfer from thalamocortical to cortical layer IV neurons to that between PNs and LHNs (1), and likewise, we further propose that these theoretical mechanisms can be applied to the thalamocortical loop and cortico-cortical feedforward projections, where spike signals propagate through multiple types of principal neurons that are different in size, morphology, ion channel expressions, etc. for each layer. Stable signal propagation in an FFN has been extensively studied in this context (3-11). However, proposed models so far were often successful only with a limited range of input signals given fixed model parameters, although some precisely tuned models can handle a diverse range of inputs (4, 6-8). In this study, we proposed a novel approach to this problem, based on information theoretic perspective, pointing at that an assumption of a single cell type in a network can result in accumulated signal distortion, whereas introducing multiple cell types with lamina-specific neuronal properties can circumvent this problem by their complementary functions. This indeed brought superior performance, exmplified by stable propagation of a dynamical spike signal. Given the prevalence of diverse cell types in many neural systems, our work presents a clear case that lamina-specific cell types are surprisingly critical to understanding network functions.

Complementarity also explains experimental observations that information encoded by an input layer appears substantially lost in the postsynaptic layer. For example, olfactory bulb output neurons simultaneously encode multiple aspects of an odor by multiplexing spike synchrony and firing rate (30), but their postsynaptic targets appear to largely filter out information in spike times due to their integrative property (31). Such phenomena naturally arise and can be explained by complementarity. In this case, postsynaptic neurons do not inherit the coding strategy of presynaptic neurons but employ a very different one. Therefore, given inputs to a presynaptic layer, postsynaptic neurons appear to respond very differently from presynaptic ones. For example, in the AL network, PNs and LHNs have different firing time courses, marked by different peak times (earlier in LHNs), primarily due to LHNs sensitively responding to correlations rather than the average variability in PN firing rates (1). Therefore, this can be misinterpreted as LHNs seemingly filtering out a substantial fraction of the information carried in a mean PN firing rate. However, we have demonstrated that on the contrary, information transfer from PNs to LHNs is in fact nearly optimal due to their complementarity. This suggests that opposing coding schemes of pre-/postsynaptic neurons, seen in experiments, can be a signature of optimal information transfer, rather than of discarding information.

Complementary transformations underlie many strategies in information theory for optimizing information transfer with a limited bandwidth, such as water-filling (32). In this study, we have demonstrated how this scheme operates in FFNs when lamina-specific neuron types have different intrinsic properties. Notably, a previous study showed that functionally different cell types within a layer can also be explained by maximization of information transmission (33). Therefore, we suggest that the commonly observed diversity of cell types in neural circuits is essential to achieve optimal information transmission.

## Materials and Methods

### Experimental procedure

Firing rates and *d*’ in Fig. 3B Bottom were based on spike times collected as described by Jeanne and Wilson (1), who kindly shared the data set. Briefly, ORNs in glomerulus DA1 of *Drosophila* antennae, expressing light-activated cation channel channelrhodopsin-2, were stimulated briefly (100 ms) by blue light emitted from an LED coupled to an optical fiber. Simultaneously, extracellular (ORNs) and patch-clamp (PNs and LHNs) recordings were performed *in vivo*. Spike times were extracted from the recording data by the custom algorithm. Although we used our custom scripts for Fig. 3B, we strictly followed the procedure in (1) to compute firing rates and *d*’ and reproduce corresponding figures faithfully.

### Model neurons

We used conductance-based model neurons based on the Morris-Lecar mechanisms (15), which are given by

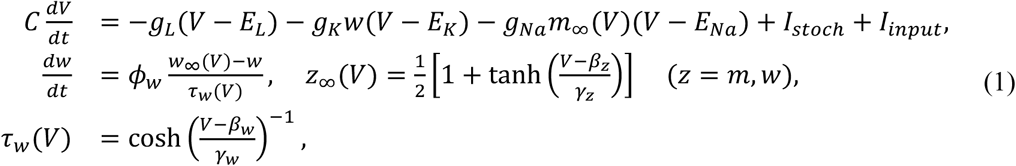

where *V* and *w* are membrane potential and a gating variable for a K^+^ channel. The first model, which we called the “integrator” neuron, had a high half-maximum voltage *β*_*w*_ while the other, “differentiator” neuron, had low *β*_*w*_. The parameters are in **Table S1**.

Stochastic current *I*_*stoch*_ represented noisy membrane potential fluctuation due to the effects that are absent from our model, such as background network inputs, an unknown noise source^1^, etc., and was given by an Ornstein-Uhlenbeck (OU) process, *dI*_*stoch*_/*dt* = -*I*_*stoch*_/*τ*_*V*_ + *σ*_*V*_ *ξ*, where *ξ* is a unit Gaussian noise, renewed each time step. *τ*_*V*_=1 ms, and *σ*_*V*_ was tuned to match experimental data in (1) (see below).

The input current *I*_*input*_ was either synaptic inputs or a common current injection to input layer neurons. Each synaptic input was conductance-based and modeled as a double exponential function: at each presynaptic spike at *t*_*s*_, the synaptic current was

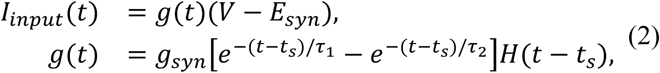

where V is the postsynaptic membrane potential, *H*(*t*) is a Heaviside function that *H*(*t*) = 1 if *t*>0 and *H*(*t*) = 0 otherwise. We used *τ*_1_=0.5 ms and *τ*_2_=4 ms, which is comparable to experimental measurments (29), and tuned *g*_*syn*_ to match experimental data as *σ*_*V*_. All other parameters are in **Table S1**. As for the current injection, see **AL network model** and **Deep FFN model** below. All simulations were constructed and run on the Brian simulator ver. 2 (34).

### AL network model

In the *Drosophila* AL network model, we used *β*_*w*_=-23 mV for ORNs to give them strong differentiator traits (35), and *β*_*w*_=5 mV and −19 mV for PN and LHN, respectively. 40 ORNs projected to each PN and 9 PNs projected to each LHN (1). The number of LHNs was 9. Each layer contained 100 replicas of these, corresponding to 100 “trials” of an experiment, which resulted in 4,000 ORNs, 900 PNs, and 900 LHNs in an entire network. We tuned synaptic conductances, *σ*_*V*_ for each layer, and peak current injection, to match experimental measurements for i) the mean spontaneous firing rates in all layers and higher cell-to-cell variability in PN firing rates (1), ii) mean peak firing rates, and iii) rate of decrease in a mean LHN firing rate. In homogeneous networks, *β*_*w*_ and *σ*_*V*_ of PNs or LHNs changed accordingly and synaptic conductances were re-tuned to match peak firing rates to the heterogeneous case.

In the simulated optogenetic activation (Fig. 3A-C), the current injected to ORNs was

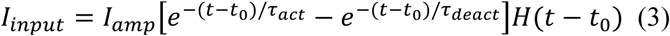

where *I*_*amp*_=45 μA/cm^2^, *τ*_*act*_=15 ms, and *τ*_*deact*_=50 ms. *t*_0_ = 200 ms is a stimulus onset. In simulations with the OU process input (Fig. 3D-E, 4D), *I*_*input*_ was again given by *dI*_*input*_/*dt* = (*μ*_*input*_*-I*_*input*_)/*τ*_*input*_+*σ*_*e*_ *ξ* where *μ*_*input*_=15 μA/cm^2^, *σ*_*input*_=7.5 μA/cm^2^, and *τ*_*input*_=5 ms. See **Table S2** for other parameters.

### Deep FFN model

All deep FFN models had 9 layers of 1,000 (5,000 in Fig. S1) differentiator or integrator neurons in the AL network model, except for the input layer composed of differentiators. Again, each neuron was randomly connected to 9 presynaptic neurons on average. Synaptic conductances and other parameters were the same as the AL network. An input layer was driven either by spikes from artificial spike generators (Fig. 4A-C, E) or by the current injection generated by an OU process (Fig. 4D). Spike generators randomly sampled in total α spike times from a normal distribution with variance σ^2^ and forced the input layer neurons to fire at the spike times, in addition to noisy spontaneous firing. The OU process case was the same as the AL network except *μ*_*input*_=25 μA/cm^2^ and *σ*_*input*_=12.5 μA/cm^2^. See **Table S3** for the other parameters.

An FFN model with feedforward inhibition (Fig. S2) had 9 layers of 4,000 excitatory neurons, and 1,000 inhibitory neurons. Each cell received 9 excitatory inputs on average from the previous layer, and each excitatory neuron also received inputs from inhibitory cells in the same layer with the same connection probability. Inhibitory cells were also based on the Morris-Lecar model (Equation 1) with *β*_*w*_ = −15 mV while different *β*_*w*_ did not cause any significant change in our conclusion. The reversal potential of inhibitory synapses was *E*_*syn*_ = −90 mV and the conductance was 200 μS/cm^2^. Also, we added a synaptic delay of 2 ms for all connections.

### Data analysis

In the AL network model case, *d*’, a measure for signal detection, was computed in the same way as in ref. (1):

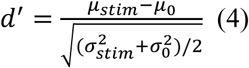

where (*μ*_stim_, *σ*_stim_) and (*μ*_0_, *σ*_0_) are the (mean, STD) of spike count at a given layer, computed with 80 ms-long overlapping temporal windows in the stimulated and non-stimulated condition, respectively. For each layer, we computed *d*’ of all the cells and plotted their median in Fig. 3B,C.

Power spectra for Fig. 3D were evaluated by applying the MATLAB function *pmtm* with a 20-ms time window on spike trains formed with 1-ms time bins. Mutual information in Fig. 3E were computed by a Gaussian channel approximation (36): We first reduced the dimensionality of a population spike trains at each layer, by using principal component analysis (PCA). Since the first PCA component was always dominating, we projected the population spike trains to this component to form a one-dimensional “population response” time series. With the Fourier transformation of the stimulus and population response, *S*(*ω*) and *R*(*ω*), we estimated a kernel *K*(*ω*) *= <R*^*^(*ω*)*S*(*ω*)>/<*R*^*^(*ω*)*R*(*ω*)>, and computed a reconstructed stimulus and noise via *S*_*r*_(*ω*) = *R*(*ω*)*K*(*ω*) and *N*(*ω*) = *S*(*ω*)-*S*_*r*_(*ω*). The mutual information per each frequency bin was then computed by

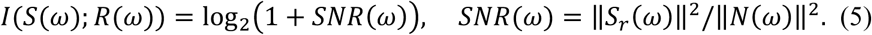

With this, we computed the information transfer (Fig. 3E) by *T*_*X*_(*ω*) = *I*(*S*(*ω*)_ORN_; *R*(*ω*)_X_)/*I*(*S*(*ω*)_ORN_; *R*(*ω*)_ORN_), where *X* is PN or LHN.

In the deep FFN, we computed (σ, α) for spikes from each layer using a custom algorithm that estimates (σ, α) in the presence of additional spontaneous firing. We first computed the baseline spontaneous firing rate *v*_0_ at each layer by averaging the firing rate obtained from the same model with no input. The firing rate curve was computed by histogramming spike times in this layer with a 0.1-ms time bin and by smoothing it with a 3-step moving average. Then, we evaluated a least-square fit of *ν*(*t*) to *ν*_*fit*_(t) = *ν*_0_ *+ ν*_1_ exp(-(*t*-*t*_*c*_)^2^/2*σ*^2^). α was estimated by counting the spikes in the [*t*_*c*_-3*σ, t*_*c*_ + 3*σ*] window. From the goodness of fit, *R*^2^ = 1 -< (*ν*(*t*)-*ν*_*fit*_(*t*))^2^>/Var[*ν*(*t*)], we evaluated the signal-to-noise ratio, *S/N = R*/(*1-R*^*2*^)^1/2^ (Fig. 4E).

The datasets generated during and/or analyzed during the current study are available from the corresponding author upon reasonable request. All the models and analysis code will be made publicly available at ModelDB (http://senselab.med.yale.edu/modeldb).

## Acknowledgements

We thank James Jeanne for great discussions and for sharing experimental data. We also thank Steven Prescott, Mario Negrello, Jihwan Myung, and Steven Aird for reading an earlier version and providing useful feedback. This work was supported by funding from the Okinawa Institute of Science and Technology Graduate University. SH was also supported by Japan Society for the Promotion of Science, KAKENHI Grant Number 15K06715.

## Author contributions

S.H. and E.D.S. conceived the research. D.H. designed and programmed computer models. D.H. and S.H. ran simulations and analyzed the data. D.H., S.H., and E.D.S. wrote the manuscript.

## Supplementary Information

**Table S1.** Parameters of the single-neuron model.

| Parameter | Value |
| --- | --- |
| $E_{Na}$ | 50 mV |
| $E_K$ | -100 mV |
| $E_L$ | -70 mV |
| $g_{Na}$ | 20 mS/cm<sup>2</sup> |
| $g_K$ | 20 mS/cm<sup>2</sup> |
| $g_L$ | 2 mS/cm<sup>2</sup> |
| $\varphi_w$ | 0.15 |
| $C$ | 2 mF/cm<sup>2</sup> |
| $\beta_w$ | -1.2 mV |
| $\gamma_m$ | 18 mV |
| $\gamma_w$ | 10 mV |
| $E_{syn, \text{ excitatory}}$ | 0 mV |
| $E_{syn, \text{ inhibitory}}$ | -90 mV |

**Table S2.** Parameters of the AL network model. η is a random number sampled from a uniform distribution ranging from 0 to 1.

| Parameter | Heterogeneous | Differentiator PN | Integrator LHN |
| --- | --- | --- | --- |
| $\beta_w$, ORN | -23 mV | -23 mV | -23 mV |
| $\beta_w$, PN | 5 mV | -19 mV | 5 mV |
| $\beta_w$, LHN | -19 mV | -19 mV | 5 mV |
| $\sigma_V$, ORN | 38 $\mu\text{A}/\text{cm}^2$ | 38 $\mu\text{A}/\text{cm}^2$ | 38 $\mu\text{A}/\text{cm}^2$ |
| $\sigma_V$, PN | $38 + 15\eta$ $\mu\text{A}/\text{cm}^2$ | 15 $\mu\text{A}/\text{cm}^2$ | $38 + 15\eta$ $\mu\text{A}/\text{cm}^2$ |
| $\sigma_V$, LHN | 15 $\mu\text{A}/\text{cm}^2$ | 15 $\mu\text{A}/\text{cm}^2$ | $38 + 15\eta$ $\mu\text{A}/\text{cm}^2$ |
| $g_{syn}$, PN | 345 $\mu\text{S}/\text{cm}^2$ | 1170 $\mu\text{S}/\text{cm}^2$ | 345 $\mu\text{S}/\text{cm}^2$ |
| $g_{syn}$, LHN | 975 $\mu\text{S}/\text{cm}^2$ | 715 $\mu\text{S}/\text{cm}^2$ | 285 $\mu\text{S}/\text{cm}^2$ |

**Table S3.** Parameters of the deep FFN model. Even and Odd represent the 2*n* and (2*n*+1)-th layer where *n*=1,2,…,5, respectively. η is a random number sampled from a uniform distribution ranging from 0 to 1.

| Parameter | Heterogeneous | All differentiator | All integrator |
| --- | --- | --- | --- |
| $\beta_w$, Input | -23 mV | -23 mV | -23 mV |
| $\beta_w$, Even | -19 mV | -19 mV | 5 mV |
| $\beta_w$, Odd | 5 mV | -19 mV | 5 mV |
| $\sigma_V$, Input | 38 $\mu\text{A}/\text{cm}^2$ | 38 $\mu\text{A}/\text{cm}^2$ | 38 $\mu\text{A}/\text{cm}^2$ |
| $\sigma_V$, Even | 15 $\mu\text{A}/\text{cm}^2$ | 15 $\mu\text{A}/\text{cm}^2$ | $38 + 15\eta$ $\mu\text{A}/\text{cm}^2$ |
| $\sigma_V$, Odd | $38 + 15\eta$ $\mu\text{A}/\text{cm}^2$ | 15 $\mu\text{A}/\text{cm}^2$ | $38 + 15\eta$ $\mu\text{A}/\text{cm}^2$ |
| $g_{syn}$, Even | 975 $\mu\text{S}/\text{cm}^2$ | 975 $\mu\text{S}/\text{cm}^2$ | 345 $\mu\text{S}/\text{cm}^2$ |
| $g_{syn}$, Odd | 345 $\mu\text{S}/\text{cm}^2$ | 975 $\mu\text{S}/\text{cm}^2$ | 345 $\mu\text{S}/\text{cm}^2$ |

**Fig. S1.**
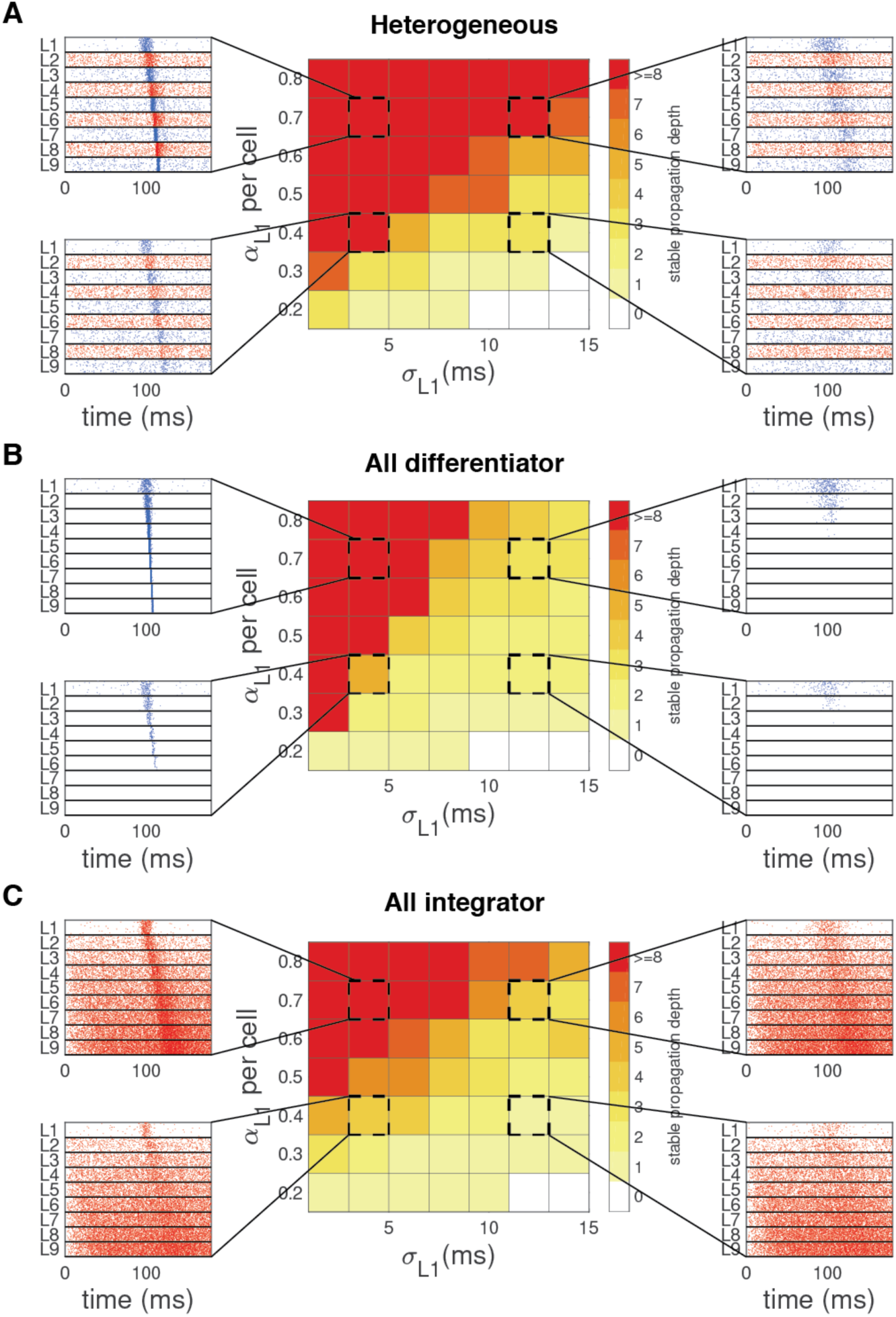
Propagation of spike signals with diverse width (σ) and number of spikes (α) in the heterogeneous (A), all-differentiator (B), and all-integrator network (C). Each network has 9 layers of 5,000 neurons (see Table S3 for parameters). Color in the middle column represents propagation depth, computed by numbers of layers (except an input layer) into which spike signals propagate. Propagation is considered stopped if the estimated α is lower than 0.05*n* or larger than 3*n* for a layer and its corresponding postsynaptic layer, where *n*=5,000 is the group size. Side insets are example raster plots for parameters marked by dotted squares in the middle, showing spikes from 10% of neurons at each layer for clarity.

**Fig. S2.**
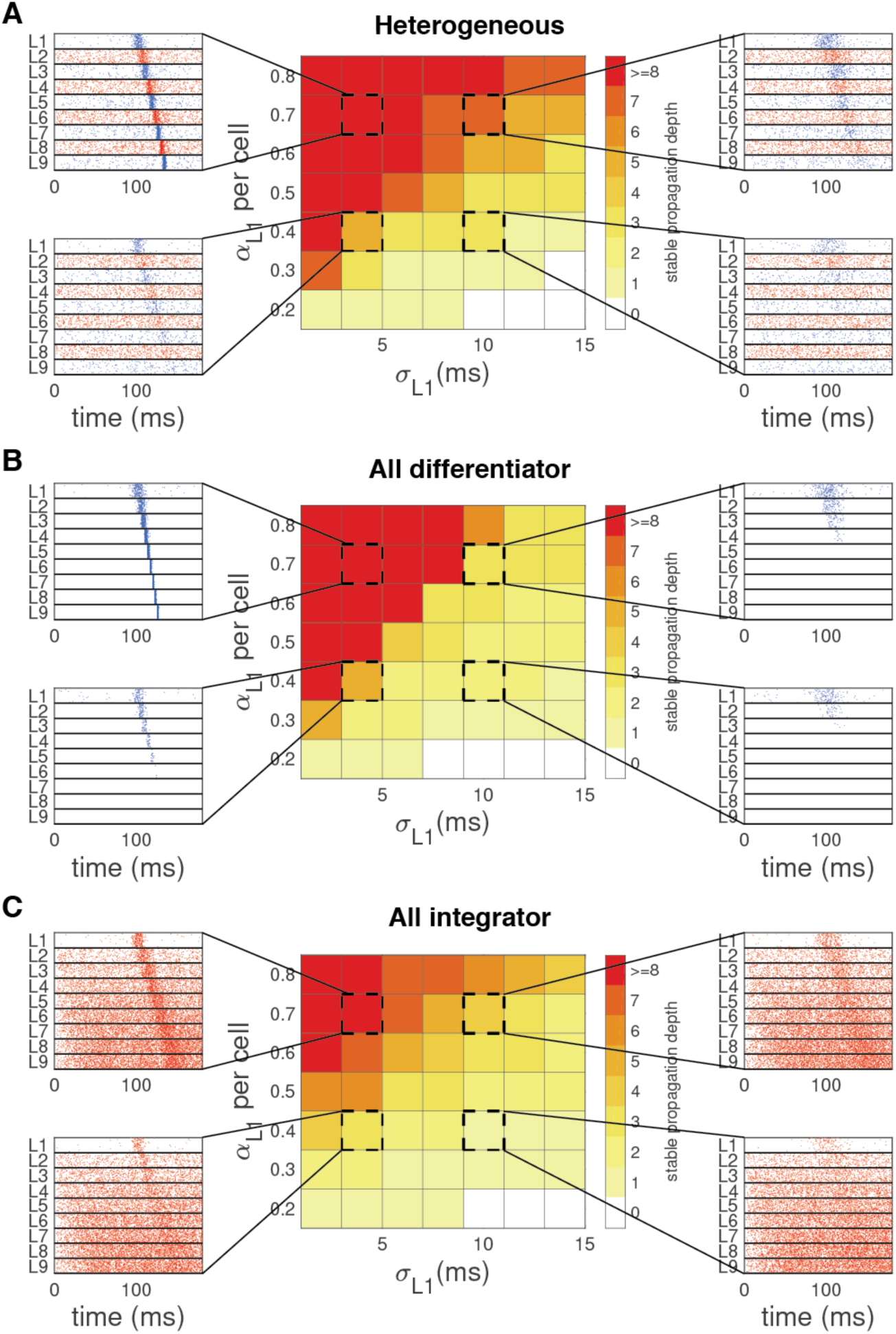
The same figures as Fig. S1, using FFN models with feedforward inhibition. Again, each network has 9 layers of 4,000 PN-like or LHN-like excitatory neurons and 1,000 inhibitory neurons that receive excitatory inputs from a previous layer and inhibit excitatory neurons in the same layer (see Methods and Table S3 for details). In all panels, we plotted spikes from 10% of excitatory neurons at each layer for clarity.

